# Periosteal and periarticular compartments house lymphatic vessels in bone

**DOI:** 10.64898/2026.04.06.716745

**Authors:** Qing Chang, Yang Shu, Hongzhi Liu, Pao-Fen Ko, Jian-Fu Chen

**Affiliations:** Center for Craniofacial Molecular Biology, University of Southern California, Los Angeles, California 90033, USA

**Keywords:** lymphatic vessel, periosteum, bone marrow, mandible, mice

## Abstract

The anatomical localization of lymphatic vessels in bone remains controversial and has led to conflicting interpretations of skeletal lymphatic function. Here we assessed lymphatic identity and localization in bone using mouse genetic labeling, tissue clearance, and three-dimensional imaging. We analyzed long bones after extensive periosteum removal and identified Vegfr3^+^ blood vessels lacking Lyve1 expression within bone marrow, whereas Vegfr3^+^Lyve1^+^ lymphatic vessels were confined to residual periosteal regions. Genetic lineage tracing using Prox1-Cre/ER;mScarlet further confirmed that lymphatic vessels are absent from long bone marrow and restricted to periosteal compartments, particularly in fibrous but not cambial layers. Extending these analyses to the mandible, we observed Vegfr3^+^Lyve1^+^ lymphatic vessels localized to periarticular soft tissues surrounding the temporomandibular joint (TMJ), while mandibular bone marrow contained only Vegfr3^+^Lyve1^−^ blood vessels and lacked Prox1 lineage-traced lymphatic vessels. Together, these findings establish that lymphatic vessels in bone are confined to periosteal and periarticular compartments and absent from bone marrow, providing a framework for interpreting lymphatic contributions to skeletal physiology and disease.

## Introduction

Historic studies suggested interstitial fluid flow in cortical bone towards lymph nodes via tracking dye flow after an injection into long bone (1, 2), which implicate the possibility of a functional lymphatic vessel system in bone. Animal models and patients with Gorham-Stout, or vanishing bone disease, exhibit lymphangiogenic bone invasion coupled with progressive osteolysis (3, 4). These studies showed the lymphatic vessels in bone under pathological conditions, while their presence during bone homeostasis is unclear. Lymphatic vessels have recently been proposed to reside within cortical bone and bone marrow under physiological conditions and to contribute directly to skeletal regeneration after injury (5). These findings have important implications for bone biology but also raise questions regarding the anatomical specificity of lymphatic vessels among different types of bones, since long bone and craniofacial bones have notable divergence in bone developmental origin, structure, and function (6).

The mandible is a specialized craniofacial bone that constitutes the movable osseous component of the temporomandibular joint (TMJ), where the mandibular condyle articulates with the temporal bone to enable jaw opening, closing, and translational movements essential for mastication, speech, and occlusion (7, 8). Like the maxilla, mandible contributes to facial architecture and oral function. However, the mandible is uniquely mobile, functioning as a load-bearing lever during dynamic mechanical activity. In contrast to long bones of the appendicular skeleton, which are primarily mesoderm-derived and form through endochondral ossification, the mandible arises largely from cranial neural crest–derived mesenchyme and develops predominantly via intramembranous ossification, with the notable exception of secondary cartilage at the condyle (9, 10). Structurally, while both long bones and the mandible contain cortical and trabecular compartments and adapt to mechanical loading, the mandible exhibits region-specific specializations reflecting its dual roles in articulation and mastication. Functionally, long bones primarily support locomotion and systemic mineral homeostasis, whereas the mandible integrates skeletal support with craniofacial biomechanics and TMJ articulation (7, 11). These developmental, structural, and functional distinctions make the mandible a compelling craniofacial model for determining whether lymphatic vessel localization patterns identified in long bone are conserved across skeletally and embryologically distinct compartments.

We recently identified lymphatic vessels in skull periosteum but not in bone marrow using genetic lineage tracing and three-dimensional (3D) imaging (12). Our result is in contrast with prior finding of lymphatic vessels in cranial bone skull (5). Here we sought to extend our lymphatic vessel studies from skull into long bone and mandible with a specific attention to the bone marrow compartment.

## Results

### Long bone marrow contains Vegfr3^+^Lyve1^−^ blood vessels but lacks lymphatic vessels

To minimize periosteal immunoreaction, we first examined long bones after extensive mechanical removal of periosteum (Fig. 1A). Our and independent groups’ studies showed that Vegfr3^+^Lyve1^-^ and Vegfr3^+^Lyve1^+^ label blood and lymphatic vessels, respectively (12–14). Using thick (100 μm) tissue sections combined with CUBIC (clear, unobstructed brain/body imaging cocktails and computational analysis) tissue clearance and confocal imaging (Fig. 1A), we detected abundant Vegfr3^+^ vessels within bone marrow that lacked Lyve1 expression in both sagittal and coronal sections of long bone (Fig. 1B-ii and 1C-iv, Video S1), suggesting their blood vessel identities. In contrast, Vegfr3^+^Lyve1^+^ double-positive lymphatic vessels were rare and restricted to residual periosteal regions (labeled by Col1a1) after extensive mechanical removal of periosteum (Fig. 1B-i and 1C-iii, Video S1). These observations indicate that long bone marrow vasculature is dominated by Vegfr3^+^Lyve1^−^ blood vessels rather than lymphatic vessels, while lymphatic vessels localize to long bone periosteum.

**Figure 1.**
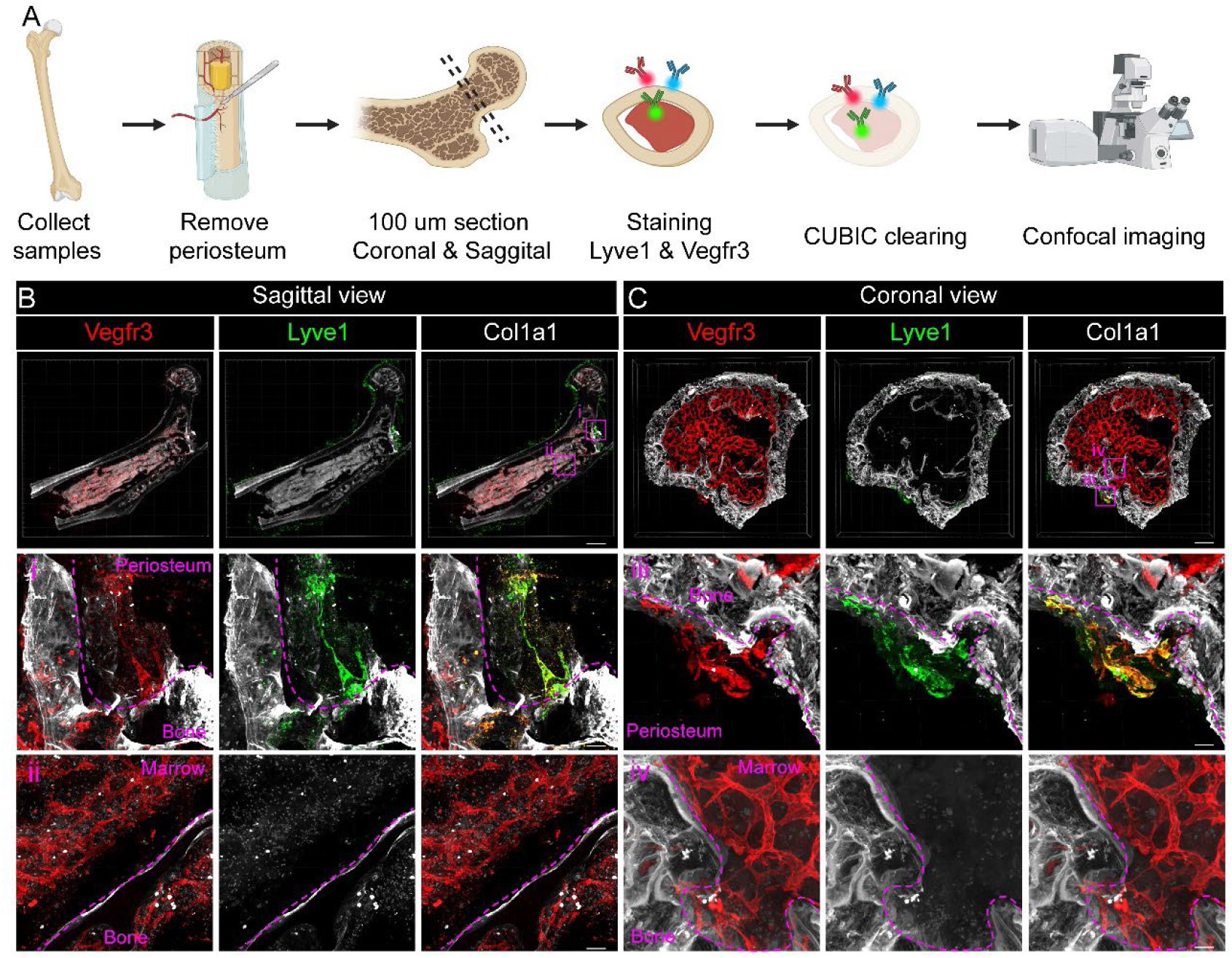
Long bone marrow contains Vegfr3^+^Lyve1^−^ blood vessels but lacks lymphatic vessels (A) Diagram of the experimental design of studying lymphatic vessels in long bone. (B, C) Confocal imaging of adult mouse long bones following extensive mechanical removal of the periosteum to selectively enrich the bone marrow compartment. Immunostaining of thick-section (100 µm) subject to CUBIC tissue clearance reveals that Vegfr3^+^Lyve1^+^ double-positive lymphatic vessels are rarely detected and are confined to residual periosteal regions in sagittal sections (B-i) and coronal sections (C-iii). There are abundant Vegfr3^+^ blood vessels (red) within bone marrow that lack Lyve1 expression (green) in both sagittal sections (B-ii) and coronal sections (C-iv). Scale bars: B, 1000 um; B-i and B-ii, 50 um; C, 200 um; C-iii and C-iv, 30 um.

### Genetic labeling reveals lymphatic vessels restricted to the fibrous out layer of periosteum and excluded from long bone marrow

To provide genetic validation, we generated *Prox1*-Cre/ER;mScarlet mice to genetically label lymphatic vessel (15). Following the femur collection, we combined with Lyve1 staining and performed CUBIC high lipids (CUBIC-HL) tissue clearance (Fig. 2A). Three-dimensional reconstruction of cleared *Prox1-CreERT2;mScarlet* femurs revealed a continuous network of Prox1 lineage-traced, Lyve1^+^ lymphatic vessels along the periosteal surface, with no detectable lymphatic structures within the intramedullary cavity (Fig. 2B, Video S2). Higher-resolution views and individual optical slices confirmed that Prox1-mScarlet^+^Lyve1^+^ vessels were restricted to periosteal and cortical-adjacent compartments, whereas the bone marrow space contained only Lyve1^−^ blood vessels and lacked Prox1 lineage labeling (Fig. 2C). The periosteum consists of an outer fibrous layer and an inner cambial layer, which can be distinguished by periostin (POSTN) expression (16). To precisely define the spatial localization of lymphatic vessels within the periosteum, we co-stained Lyve1 and POSTN in Prox1-mScarlet long bone. High-magnification views demonstrate that Prox1-mScarlet^+^Lyve1^+^ lymphatic vessels are confined to the fibrous outer layer and are excluded from the POSTN^+^ cambial inner layer (Fig. 2D). These data demonstrate that long bone lymphatic vessels are specifically restricted to the periosteal fibrous layer and are excluded from both the cambial layer and bone marrow (Fig. 2A).

**Figure 2.**
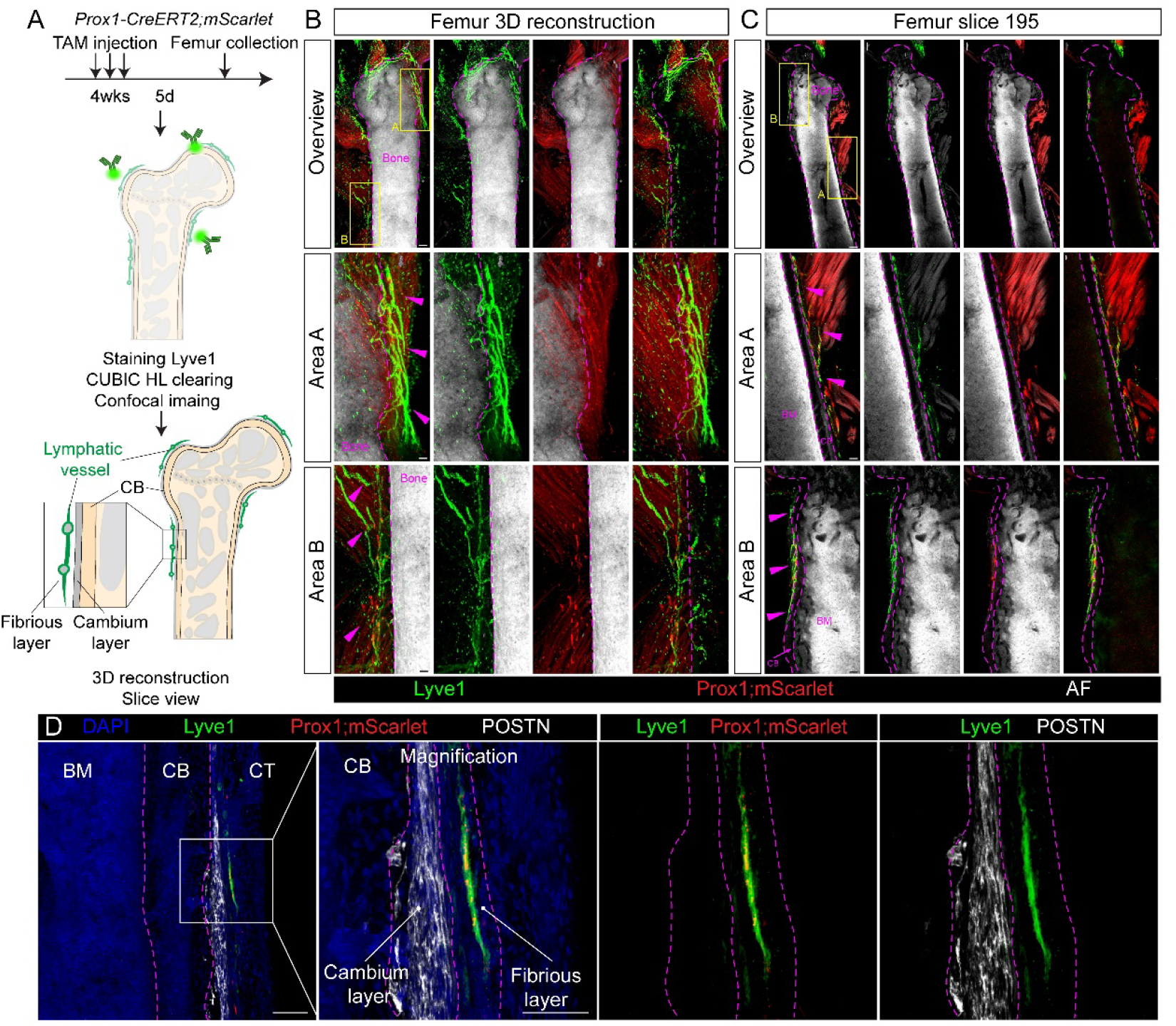
Prox1 genetic labeling reveals lymphatic vessels restricted to the periosteum and excluded from long bone marrow (A)Experimental schematic illustrating tamoxifen (TAM) induction of *Prox1-CreERT2;mScarlet* mice, followed by femur collection, Lyve1 immunostaining, CUBIC-HL tissue clearing, confocal imaging and three-dimensional reconstruction. (B)Three-dimensional reconstruction of cleared femurs shows Prox1-mScarlet^+^ (red) Lyve1^+^ (green) lymphatic vessels forming a continuous network along the outer cortical surface and periosteal compartment (outlined by dashed magenta lines). Higher-magnification views of boxed regions (Areas A and B) reveal Prox1 lineage-traced lymphatic vessels closely apposed to the periosteum (arrowheads), while the intramedullary space lacks Prox1-mScarlet^+^Lyve1^+^ structures. (C)Representative single optical slice from the same femur confirms the three-dimensional observations, showing Prox1-mScarlet^+^Lyve1^+^ lymphatic vessels confined to periosteal and cortical-adjacent regions, whereas the bone marrow cavity contains no Prox1 lineage-traced lymphatics. (D)Representative immunofluorescence images of long bone showing DAPI (blue), Lyve1 (green), Prox1-mScarlet (red), and periostin (POSTN, white). POSTN labels the cambial inner layer of the periosteum adjacent to cortical bone (CB), distinguishing it from the outer fibrous layer. Prox1-mScarlet^+^Lyve1^+^ lymphatic vessels are localized to the fibrous layer and are excluded from the POSTN^+^ cambial layer and bone marrow (BM). Dashed lines indicate periosteal boundaries. Boxed area is shown at higher magnification. CB, cortical bone; CT, connective tissue.

### Lymphatic vessels localize to periarticular soft tissues but not bone marrow in mandible

We next extended this analysis to an independent craniofacial skeletal site, the mandible, focusing on the temporomandibular joint (TMJ) (Fig. 3A). We recently identified lymphatic vascular system in the TMJ synovium and defined its function in arthritis and pain (17). However, the detailed anatomical locations of lymphatic vessels in non-synovial tissues such as condyle bone have not been documented. We used well-established Complete Freund’s Adjuvant (CFA)-induced inflammatory TMJ arthritis mouse models with robust lymphangigogenesis to maximize lymphatic vessel detection (18). CUBIC tissue clearance of 100 µm-thick mouse TMJ sections again revealed a clear compartmentalization: Vegfr3^+^Lyve1^+^ lymphatic vessels formed continuous, tube-like structures in periarticular soft tissues, including synovium, muscle, retrodiscal tissue, periosteum, and joint capsule (Fig. 3B, upper panels, Video S3). Mandible zoom in analysis showed that mandibular bone marrow contained only Vegfr3^+^Lyve1^−^ blood vessels (Fig. 3B-i, white arrowheads). In contrast, adjacent mandible periosteum exhibited Vegfr3^+^Lyve1^+^ lymphatic vessels (Fig. 3B-ii, purple arrowheads). Thus, across both long bone and mandibular models, lymphatic vessels localized to periosteal and periarticular regions and outside the bone marrow compartment.

**Figure 3.**
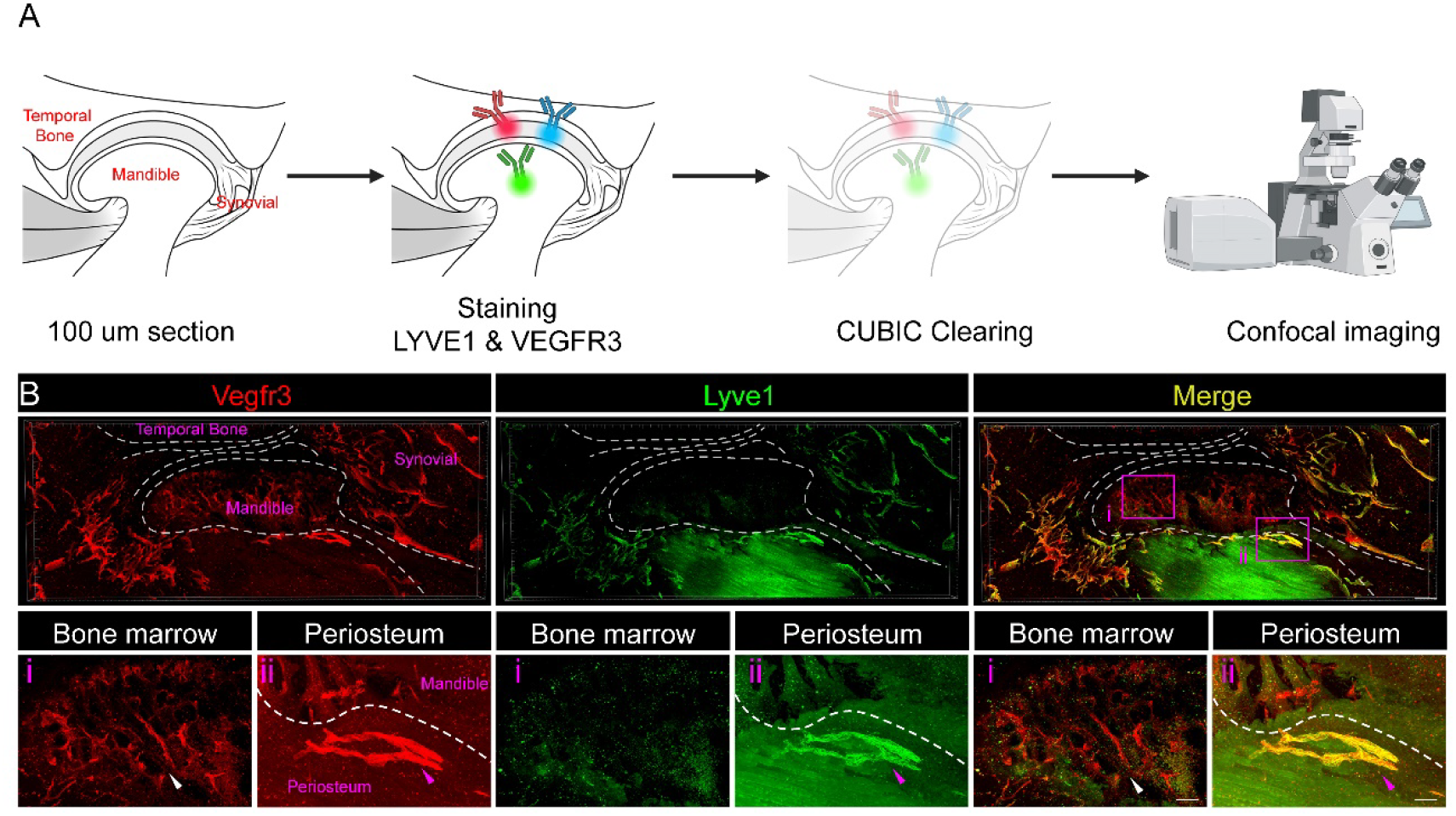
Lymphatic vessels localize to periarticular soft tissues but not bone marrow in mandible (A)Diagram of the experimental design of studying lymphatic vessels in TMJ mandible. (B)Confocal imaging of adult mouse TMJ after intra-articular injection of CFA for two weeks to induce arthritis. TMJ sections (100 µm) are subject to immunostaining for Vegfr3 and Lyve1 followed by CUBIC tissue clearance. Vegfr3^+^Lyve1^+^ lymphatic vessels form continuous, tube-like structures in periarticular regions including synovial tissue, muscle, and joint capsule surrounding the mandible. Note that the mandibular bone marrow contains Vegfr3^+^Lyve1^−^ vessels (i) and lacks Vegfr3^+^Lyve1^+^ lymphatic structures. White arrowheads indicate Vegfr3^+^ blood vessels (red) that lack Lyve 1 (green). Purple arrowheads indicate Vegfr3^+^Lyve1^+^ lymphatic vessels at the periosteum (ii). These data demonstrate distinct compartmentalization of lymphatic vessels versus blood vessels in the mandibular joint region. Scale bars: B, 200 um; B-i and B-ii, 50 um.

### Genetic tracing confirms periarticular presence of lymphatic vessels and their absence in mandibular bone marrow

Lastly, we generated *Prox1*-Cre/ER;mScarlet mice to genetically label lymphatic vessel in TMJ. Following the microdissection of the whole TMJ, we combined with Lyve1 staining and performed CUBIC high lipids (CUBIC-HL) tissue clearance, followed by confocal imaging and reconstruction (Fig. 4A). Prox1^+^Lyve1^+^ lymphatic vessels were readily detected in periarticular tissues at synovial regions, muscle, and periosteal region (Fig. 4B). These vessels form tube-like lymphatic network surrounding the synovial capsule but remain spatially restricted to peri-articular soft tissues (Fig. 4B, Video S4). We generated a movie to sequentially capture 2D slice views (medial-lateral) extracted from the 3D TMJ reconstruction (Video S4), in which autofluorescence (AF, grayscale) outlines major TMJ anatomical structures. As showing in representative sections (Fig. 4C-i-iv), Prox1^+^Lyve1^+^ double-positive lymphatic tubes are consistently detected along the synovial tissues and periosteum of the mandible (purple arrowheads). However, there are no lymphatic structures observed within bone marrow. This genetic evidence confirms that lymphatic vessels are excluded from mandibular bone marrow and validates the marker-based findings of their presence at periarticular compartments.

**Figure 4.**
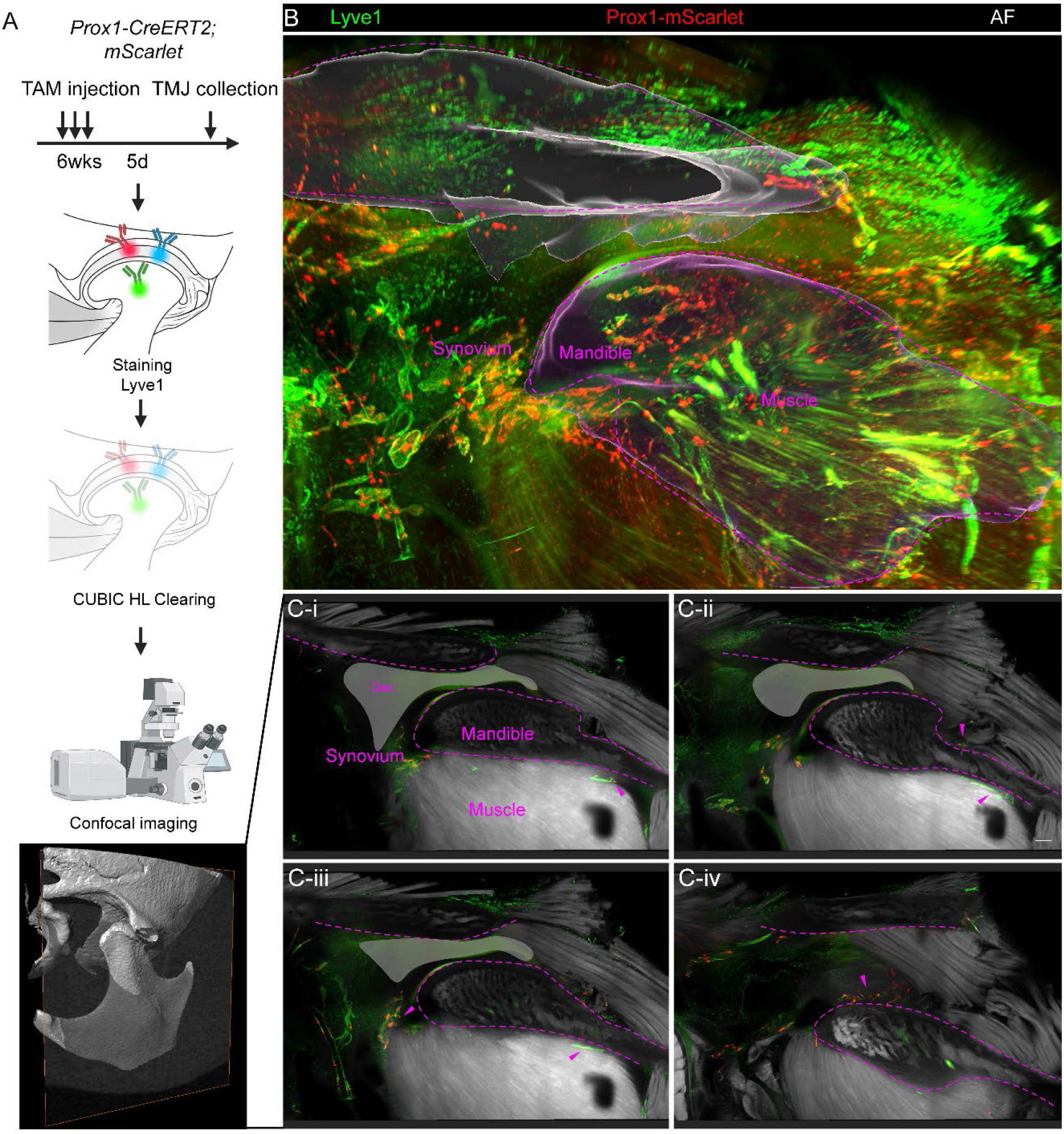
Genetic labeling confirms periarticular presence of lymphatic vessels and their absence in mandibular bone marrow (A)Schematic overview of the experimental workflow. *Prox1-CreERT2;mScarlet* mice were administered tamoxifen at 6 weeks of age, and TMJ tissues were collected 5 days after induction. Whole TMJ samples were permeabilized using the CUBIC-HL reagent, followed by immunostaining with anti-Lyve1 (Alexa Fluor 488). Samples were subsequently cleared using the CUBIC-HL protocol and imaged by confocal microscopy to obtain a complete 3D reconstruction of the joint. Whole-mount 3D rendering of the cleared TMJ, showing Lyve1^+^ lymphatic vessels (green) and endogenous *Prox1*-mScarlet fluorescence (red). Prox1^+^/Lyve1^+^ double-positive tubular structures are prominently localized within the TMJ synovium and muscle but absent from bone marrow. These vessels form a well-organized lymphatic network surrounding the synovial capsule and remain spatially restricted to peri-articular soft tissues. Scale bars: B, 300 um. (C, i–iv) Sequential 2D slice views (medial → lateral) extracted from the 3D TMJ reconstruction. Autofluorescence (AF, grayscale) outlines major anatomical structures. Each panel highlights the spatial relationship between lymphatic vessels and joint structures, including the mandibular bone, synovium, muscle, and articular disc. Across sections C-i to C-iv, Prox1^+^/Lyve1^+^ double-positive lymphatic tubes are consistently detected along the periosteum of the mandible, but no lymphatic structures are observed within bone marrow. Scale bars: C-i-iv, 100 um.

## Discussion

The anatomical presence of lymphatic vessels within bone has remained controversial, with prior studies proposing lymphatic networks within cortical bone and marrow that contribute to skeletal homeostasis and repair injury (5). In this study, we provide convergent evidence using marker-based identification, genetic lineage tracing, and three-dimensional imaging that lymphatic vessels are excluded from bone marrow and instead are restricted to periosteal and periarticular compartments across both long bone and craniofacial mandible. These findings establish a revised anatomical framework for skeletal lymphatics and clarify inconsistencies in the field.

A key advance of this study is the resolution of ambiguity arising from marker overlap between blood and lymphatic endothelium. Vegfr3 expression has been widely used to identify lymphatic vessels but is also expressed in subsets of blood vessels, particularly under developmental or remodeling conditions (13, 19). By combining Vegfr3 with Lyve1 and validating findings using Prox1 lineage tracing, we demonstrate that Vegfr3^+^Lyve1^−^ vessels within bone marrow represent blood vasculature rather than lymphatic structures. This distinction is critical, as prior reports of “lymphatics” in bone marrow may have been confounded by reliance on single markers or limited spatial resolution (5). Our use of thick-section clearing and 3D reconstruction further minimizes the risk of misinterpreting periosteal contamination as intramedullary localization.

Our data consistently localize lymphatic vessels to the fibrous layer of the periosteum, excluding the cambial layer and marrow cavity. This spatial restriction suggests that lymphatic vessels are anatomically positioned to regulate interstitial fluid drainage, immune surveillance, and molecular transport at the bone surface, rather than within the mineralized or hematopoietic compartments.

The periosteum is highly vascularized and innervated and plays central roles in bone repair and mechanosensation (20, 21). Localization of lymphatics to the fibrous layer raises the possibility that lymphatic vessels coordinate with periosteal blood vessels, sensory nerves, and stromal cells to modulate inflammatory responses and tissue remodeling at the bone interface. Notably, the exclusion of lymphatics from the cambial layer, which harbors osteoprogenitors (16), begs the questions how lymphatic vasculature interacts with skeletal progenitor cells and osteogenesis for bone regeneration.

Extending these findings to the mandible and TMJ provides important insight into craniofacial skeletal biology. Despite fundamental differences in embryonic origin, ossification mode, and mechanical function between long bones and the mandible (7, 22), we observed a conserved pattern in which lymphatic vessels are absent from bone marrow but enriched in periarticular and periosteal soft tissues. In the TMJ, lymphatic vessels form networks within synovium, retrodiscal tissue, and surrounding musculature (17), consistent with their established roles in joint fluid homeostasis and inflammation. Their absence from mandibular bone marrow, even under conditions of robust inflammation and lymphangiogenesis (23, 24), underscores the strength of compartmental restriction and argues against inducible lymphatic invasion into bone marrow under these conditions.

These findings have several implications for skeletal physiology and disease. First, they suggest that previously observed tracer movement from bone to lymph nodes may occur via periosteal or perivascular pathways, rather than through intramedullary lymphatic vessels (1, 2). Second, they refine our understanding of pathological conditions such as Gorham-Stout disease (3, 4), where lymphatic invasion into bone likely represents a disease-associated breach of anatomical boundaries rather than an exaggeration of a physiological baseline. Third, they provide a framework for interpreting recent studies linking lymphatics to bone regeneration (25, 26), suggesting that lymphatic effects may be mediated through periosteal niches or adjacent soft tissues rather than direct interaction with marrow-resident cells.

Overall, this work underscores the necessity of careful spatial separation of periosteum and bone marrow, multiple marker combination, and genetic lineage tracing when assessing lymphatic vessel identity and localization in mineralized tissues. Our findings refine the anatomical interpretation of bone-associated lymphatic signals reported previously(5). Together with our skull studies (12), these data indicate that lymphatic vessels in bone localize to periosteal and periarticular compartments rather than residing within bone marrow, providing a clarified framework for lymphatic functions in skeletal biology and disease.

### Limitations of the study

while our approaches provide high spatial resolution and genetic specificity, they focus on steady-state conditions and a single inflammatory model in the TMJ. It remains possible that more severe injury models, aging, or tumor-associated remodeling could alter lymphatic distribution. In addition, while Prox1 lineage tracing is a robust marker of lymphatic identity, transient or noncanonical lymphatic-like states cannot be fully excluded. Future studies integrating functional assays, live imaging, and additional lineage tools will be important to further define lymphatic behavior in skeletal contexts.

## Methods

### Sex as a biological variable

Sex was not considered as a biological variable in this study. All experiments were conducted using mice of a single sex (female). This approach was adopted to maintain experimental consistency and reduce variability across imaging and genetic analyses. The anatomical and cellular features investigated, including lymphatic vessel identity and spatial localization in bone and periosteal compartments, are not expected to be sex-dependent. Therefore, the findings are anticipated to be broadly applicable across sexes; however, this was not directly tested in the present study.

### Animals

All animal procedures were performed in compliance with protocols approved by the Institutional Animal Care and Use Committee (IACUC) at the University of Southern California. C57BL/6J mice are from the Jackson Laboratory. Prox1-CreERT2 and Ci1H11-CAG-LSL-mScarlet reporter mice were gift from Drs. Young-Kwon Hong and Hu Zhao, respectively. Cre-dependent reporter expression was induced by daily intraperitoneal administration of tamoxifen (Sigma-Aldrich, T5648; 20 mg/ml in corn oil) at a dose of 1.5 mg per 10 g of body weight.

### Cryosectioning and CUBIC clearing

Mice were transcardially perfused through the left ventricle with ice-cold PBS followed by 4% paraformaldehyde (PFA) after opening the right atrium. Skulls were then collected, post-fixed overnight, and decalcified in 20% EDTA at 4°C for 3 days. For TMJ samples, decalcification was carried out for 10 days, whereas long bone samples were decalcified for 7 days. For TMJ and lone bone whole mount tissue clearing, the samples were treated with CUBIC-L (T3740, TCI) for 3 days and blocked with 5% donkey serum for 1 day. Then the sample underwent immunostaining with anti-LYVE1 (goat polyclonal, AF2125, R&D) for 7 days and secondary antibody for 7 days before treating with CUBIC-R (T3741, TCI). For cryosection preparation, decalcified tissues were dehydrated in 30% sucrose and embedded in OCT. Samples were subsequently sectioned at 100 µm using a Leica cryostat microtome. The thick (100 µm) sections were treated overnight with CUBIC-L before immunostaining. Primary antibodies used for immunostaining were anti-LYVE1(goat polyclonal, AF2125, R&D), anti-Lyve-1 (rabbit polyclonal, ab33682, Abcam), anti-VEGFR3 (goat polyclonal, AF743, R&D), and anti-Periostin (goat polyclonal, AF2955, R&D). Sections were incubated with primary antibodies for 2 days at 37°C, and nuclei were counterstained with DAPI (Invitrogen). Appropriate secondary antibodies were applied according to the host species of the primary antibodies. After staining, sections were cleared and mounted using CUBIC-R. Mounted skull specimens were placed in confocal glass-bottom dishes (Cat. 801002, NEST Inc.) for imaging. All antibodies used in this study were validated by the manufacturers for the relevant species and experimental applications.

### Confocal imaging and analysis

Prepared samples were imaged using a Leica STELLARIS 5 confocal microscope. Whole-skull imaging was performed with a 10×/0.3 NA objective lens with a 1000 µm working distance. Following whole-skull imaging, selected skull fragments were dissected for higher-resolution imaging with a 63×/1.4 NA objective lens with a 200 µm working distance. Image acquisition was carried out using LAS X Life Science Microscope Software (version 1.4.6). The system was equipped with fixed laser lines at 405, 488, 561, and 633 nm. High-magnification scans were acquired at 63×/1.40 NA with a z-step size of 0.8 µm using continuous scanning at 488, 561, and 633 nm. Raw confocal data were gamma-adjusted to enhance visualization of the images. File format conversion was performed in ImageJ, and Bitplane Imaris was used for three-dimensional reconstruction, manual 3D annotation, and movie generation.

### Study approval

All animal experiments were performed in accordance with protocols approved by the Institutional Animal Care and Use Committee (IACUC) at the University of Southern California. All procedures were conducted in compliance with institutional guidelines for the care and use of laboratory animals. No human participants were involved in this study; therefore, written informed consent was not applicable. No patient photographs were included in this study.

## Acknowledgments

We thank Chen laboratory colleagues for stimulating discussions. We thank Drs. Young-Kwon Hong and Hu Zhao for providing *Prox1*-Cre^ERT2^ and Ci1H11-CAG-LSL-mScarlet mice.

## Author Contributions

Q.C., Y.S., H-Z.L., and P-F.K. conceived and performed all experiments.

J-F.C designed and interpreted the experiments and wrote the manuscript.

## Financial Statement

The authors declare there are no competing financial interests that might be perceived as affecting the objectivity of these studies.

