## Supplemental material for "Periosteal and periarticular compartments house lymphatic vessels in bone"

**Video S1.** Video analysis of confocal imaging of CUBIC-cleared 100 μm-thick mouse long bone sections.

**Video S2.** Video analysis of confocal imaging of CUBIC-HL-cleared femur with genetic labeling of lymphatic vessels (*Prox1*-mScarlet, red) coupled with Lyve1 staining (green). Genetically labeled lymphatic vessels (*Prox1*-Cre–driven mScarlet$^+$, Lyve1$^+$) form continuous, tube-like structures in periosteal regions. In contrast, no Prox1$^+$Lyve1$^+$ lymphatic vessels are detected within bone marrow.

**Video S3.** Video analysis of confocal imaging of CUBIC-cleared 100 μm-thick mouse TMJ sections.

**Video S4.** Video analysis of confocal imaging of CUBIC-HL-cleared whole mouse TMJ with genetic labeling of lymphatic vessels (*Prox1*-mScarlet, red) coupled with Lyve1 staining (green). Genetically labeled lymphatic vessels (*Prox1*-Cre–driven mScarlet$^+$, Lyve1$^+$) form continuous, tube-like structures in periarticular regions, including synovial tissues, adjacent muscle, periosteum, and joint capsule surrounding the mandibular condyle. In contrast, no Prox1$^+$ Lyve1$^+$ lymphatic vessels are detected within mandibular bone marrow. Depth-resolved rendering and orthogonal views confirm the spatial restriction of lymphatic vessels to periarticular soft tissues and their exclusion from mineralized bone compartments.
